# Loss of PKCα increases arterial medial calcification in a uremic mouse model of chronic kidney disease

**DOI:** 10.1101/2020.05.20.097642

**Authors:** Samantha J Borland, Cecilia Facchi, Julia Behnsen, Antony Adamson, Neil E Humphreys, Philip J Withers, Michael J Sherratt, Sheila E Francis, Keith Brennan, Nick Ashton, Ann E Canfield

## Abstract

Arterial medial calcification is an independent risk factor for mortality in chronic kidney disease. We previously reported that knock-down of PKCα expression increases high phosphate-induced mineral deposition by vascular smooth muscle cells *in vitro*. This new study tests the hypothesis that PKCα regulates uremia-induced medial calcification *in vivo*. Female wild-type and PKCα^−/−^ mice underwent a two-stage subtotal nephrectomy and were fed a high phosphate diet for 8 weeks. X-ray micro computed tomography demonstrated that uremia-induced medial calcification was increased in the abdominal aorta and aortic arch of PKCα^−/−^ mice compared to wild-types. Blood urea nitrogen was also increased in PKCα^−/−^ mice compared to wild-types; there was no correlation between blood urea nitrogen and calcification in PKCα^−/−^ mice. Phosphorylated SMAD2 immunostaining was detected in calcified aortic arches from uremic PKCα^−/−^ mice; the osteogenic marker Runx2 was also detected in these areas. No phosphorylated SMAD2 staining were detected in calcified arches from uremic wild-types. PKCα knock-down increased TGF-β1-induced SMAD2 phosphorylation in vascular smooth muscle cells *in vitro*, whereas the PKCα activator prostratin decreased SMAD2 phosphorylation. In conclusion, loss of PKCα increases uremia-induced medial calcification. The PKCα/TGF-β signaling axis could therefore represent a new therapeutic target for arterial medial calcification in chronic kidney disease.

## Introduction

Chronic kidney disease (CKD) is a major healthcare burden, with cardiovascular disease a leading cause of death among end-stage renal disease patients^1^. Vascular calcification is a hallmark feature of CKD, and calcification levels are positively correlated with increased morbidity and mortality in these patients^2,3^. Vascular calcification is an active, cell-regulated process resulting in the formation of mineralized tissue, bone and/or cartilage within the blood vessel wall^4^, occurring either within the medial layer of the vessel wall or in the intimal layer in association with atherosclerosis. While CKD patients can develop both types of vascular calcification, calcification of the medial layer is more specific to CKD and is the exclusive form of vascular calcification observed in pediatric CKD patients^5^. Vascular calcification reduces arterial (including aortic) elasticity, leading to an increase in pulse wave velocity, development of left ventricular hypertrophy, reduced coronary perfusion, and myocardial infarction and failure^6^. These cardiovascular complications are the main causes of mortality in patients with CKD. Vascular calcification is also associated with a higher mortality risk in CKD patients following kidney transplantation, and may be a predictor of poor graft outcomes^7^. As existing treatments for vascular calcification in CKD patients are limited, there is an urgent need to identify new therapeutic targets for this disease.

Vascular calcification shares many features with bone formation^8–10^. The phenotypic modulation of a subset of vascular smooth muscle cells (VSMCs) has been strongly implicated in this disease process as these cells can differentiate into osteo/chondrogenic-like cells both *in vitro*^11–16^ and *in vivo*^17,18^ in response to specific stimuli, resulting in the deposition of a mineralized matrix in the vessel wall. Degradation of elastin in the blood vessel wall has also been shown to occur in uremia and CKD^19^, leading to the over-expression of transforming growth factor-β (TGF-β) which promotes VSMC osteogenic differentiation and matrix mineralisation^20,21^.

Protein kinase Cα (PKCα) is a serine/threonine protein kinase that is ubiquitously expressed *in vivo*^22^. It was previously proposed that PKCα inhibitors may improve renal function in patients with diabetic nephropathy^23^. However, we recently discovered that biochemical inhibition of PKCα activity using Gö6976, or knocking-down PKCα expression using small interfering RNA (siRNA), increases high phosphate-induced mineral deposition by vascular smooth muscle cells (VSMCs) *in vitro*^20^; suggesting that vascular calcification and its devastating consequences could be increased if renal disease patients are treated with PKCα inhibitors. However, whether loss of PKCα increases calcification in the inherently more complex environment *in vivo* was unknown.

Therefore, this study aimed to determine whether loss of PKCα affects uremia-induced arterial medial calcification *in vivo* using the murine sub-total nephrectomy and high phosphate diet model^24–26^. Using novel X-ray micro computed tomography (μCT) approaches, we demonstrate that uremia-induced arterial medial calcification is increased significantly in PKCα^−/−^ mice compared to wild-type controls. Furthermore, both *in vitro* and *in vivo* studies suggest this increase in calcification is mediated by up-regulated TGF-β/SMAD2 signaling in PKCα knock-out VSMCs.

## Results

### Generation of PKCα^−/−^ mice using CRISPR/Cas9

For studies on uremia-induced arterial medial calcification, it is essential that mice are on a DBA/2 background to facilitate rapid mineralisation before mice enter into end-stage renal disease^25–27^. Mice are typically backcrossed for at least 5 generations onto the DBA/2 strain; however, this approach can be timeconsuming due to the poor breeding patterns encountered by this strain. Therefore, we generated PKCα^−/−^ mice on the DBA/2 background using CRISPR/Cas9 (Figure 1A). Mice harbouring InDel-containing alleles were identified by size change from wild-type amplicon size (230bp) (Figure 1B). Three mice were taken forward for Sanger sequencing and changes predicted to lead to frameshift mutations in the exon, and thus gene knock-out, were identified (Figure 1C). Mouse founder 1 was bred with wild-type DBA/2 to establish a colony and bred to homozygosity. Knock-out of the PKCα gene was confirmed by immunoblotting (Figure 1D).

**Figure 1.**
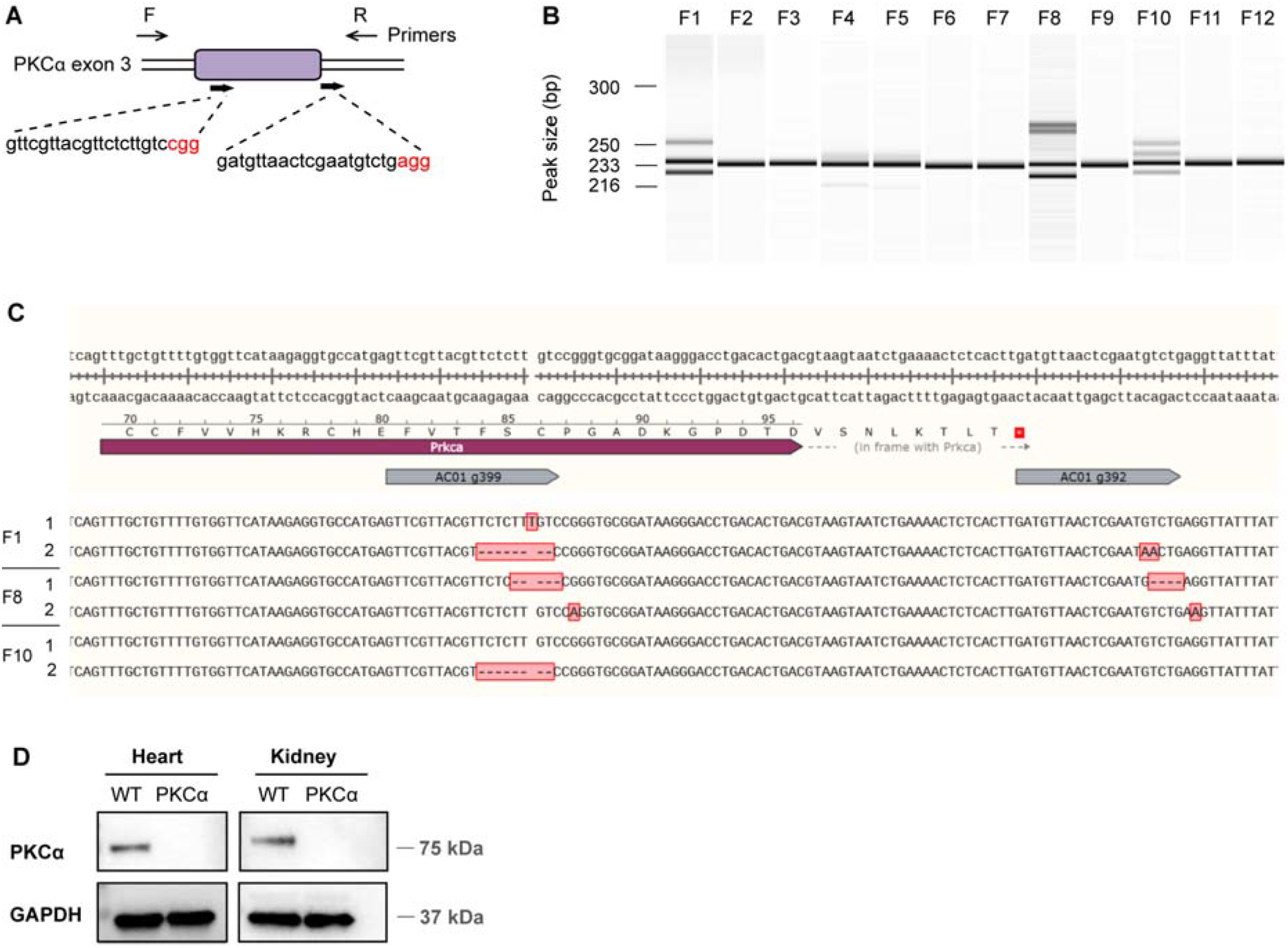
Generation of PKCα^−/−^ mice. (**A**) CRISPR targeting design of PKCα exon 3, sgRNA sequences (blue) and PAM sites (red) indicated. (**B**) PCR genotyping of founders 1-12 (F1-F12). Multi-bands were potential InDel/knock-out alleles. (**C**) Sequencing of founders 1, 8 and 10. Two different sequences were detected for each pup and are displayed aligned to wild-type (WT) sequence; changes to WT are highlighted in red shaded boxes and summarised in Supplemental Table 1. (**D**) Heart and kidney tissue lysates from wild-type and PKCα^−/−^ mice were immunoblotted for PKCα. GAPDH was used as a loading control.

### PKCα^−/−^ mice exhibit greater uremia following sub-total nephrectomy and high phosphate diet

To determine the role of PKCα in uremia-induced arterial medial calcification, we used the two-stage subtotal nephrectomy and high phosphate diet model^24–26^. Female mice were used as our pilot studies showed that they were more resistant to acute renal failure following the uni-nephrectomy in the second surgical procedure compared with males (results not shown). By the end of the experimental protocol, 77% of female wild-type and 80% of female PKCα^−/−^ mice survived (Figure 2A). There was no significant difference between the survival rates of female wild-type and PKCα^−/−^ mice, suggesting the observed mortality rate is due to consequences of the sub-total nephrectomy itself rather than loss of PKCα. Indeed, 80% of the animal losses occurred within 24 hours of the second surgery, the point at which the risk of acute renal failure is highest. There was no difference in body mass between wild-type and PKCα^−/−^ mice for the duration of the protocol (results not shown).

**Figure 2.**
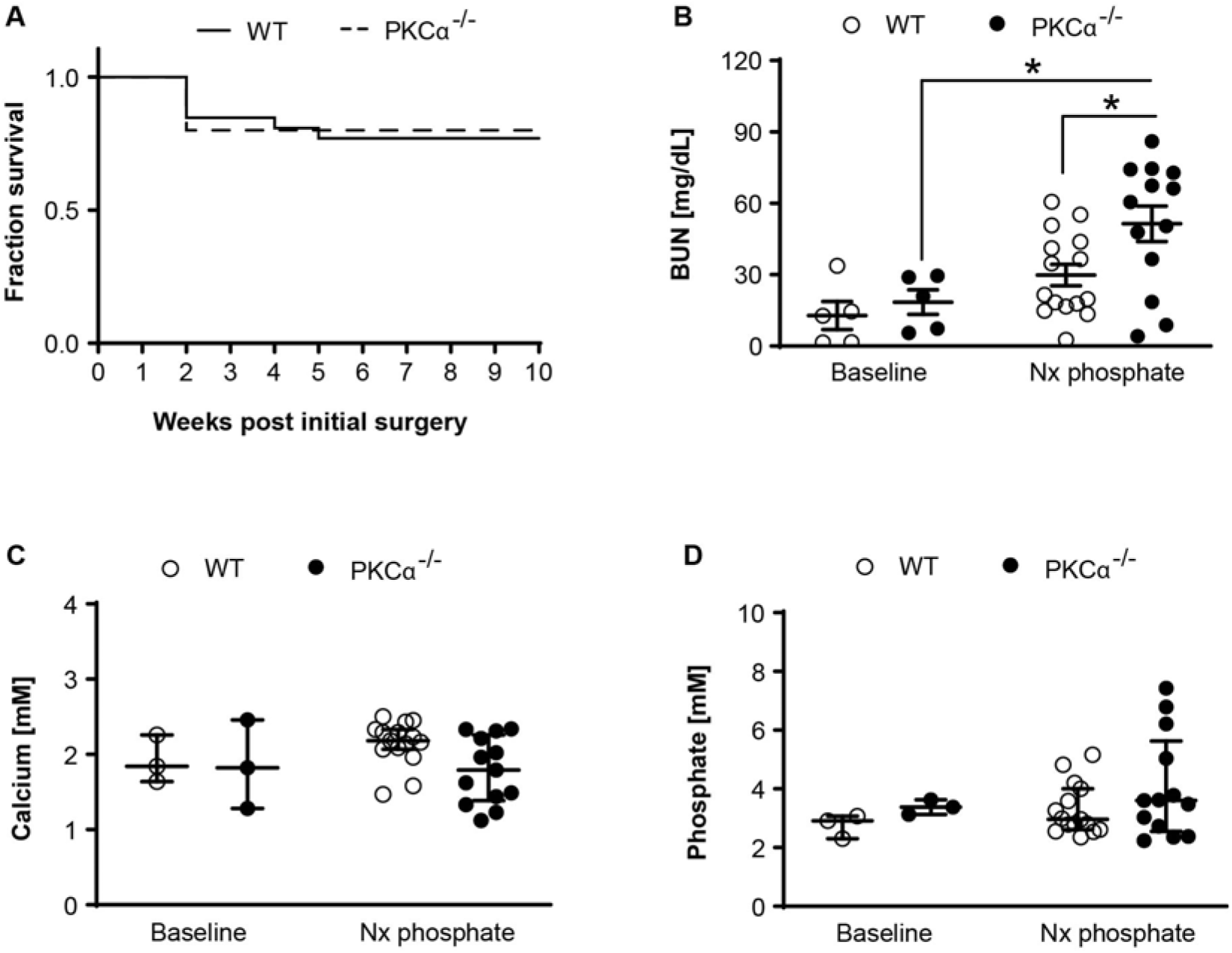
Survival and serum biochemistry for 5/6 nephrectomy and high phosphate diet fed wildtype and PKCα^−/−^ mice. (**A**) Kaplan-Meier survival curve of wild-type (WT) and PKCα^−/−^ mice following 5/6 nephrectomy and high phosphate diet feeding. Gehan-Breslow-Wilcoxon test (P=0.9018, not significant). Plasma (**B**) BUN, (**C**) calcium and (**D**) phosphate values for WT and PKCα^−/−^ mice at baseline (WT n=3-5; PKCα^−/−^ n=3-5) and the end of study following 5/6 nephrectomy and high phosphate diet feeding (Nx phosphate) (WT n=15; PKCα^−/−^ n=13). (**B**) Data expressed as mean ± SEM and were analyzed using a 2-way ANOVA and Tukey post-hoc comparisons. (**C, D**) Data expressed as median ± interquartile ranges and were analyzed using a Kruskall-Wallis test. *P<0.05.

Uremia is a surrogate marker of renal function^8^; thus plasma BUN levels were measured at the end of the experimental protocol and compared to age-matched mice at the start of the study. BUN levels were raised in both wild-type and PKCα^−/−^ mice when compared to baseline (Figure 2B), confirming induction of uremia; this increase was significant in PKCα^−/−^ mice (2.4-fold increase, P<0.05; Figure 2B). By the end of the experimental protocol, BUN levels were increased significantly in PKCα^−/−^ mice when compared to wild-type controls (1.5-fold increase, P<0.05; Figure 2B). There was no correlation between BUN levels and the degree of renal mass reduction in either genotype (results not shown).

Following sub-total nephrectomy and high phosphate diet feeding for 8 weeks, there was no significant change in plasma calcium (Figure 2C) and phosphate (Figure 2D) levels in either wild-type or PKCα^−/−^ mice when compared to baseline. Calcium (Figure 2C) and phosphate (Figure 2D) levels were also similar between wild-type and PKCα^−/−^ mice at the end of the experimental protocol.

### Arterial medial calcification is increased in PKCα^−/−^ mice

Compared with conventional histological approaches, μCT can visualise and quantify calcification throughout a whole blood vessel in 3-D without the potential for artefacts caused by mechanical sectioning^28, 29^. Therefore, μCT was utilised in this study to quantify calcification in the aortic arch and abdominal aorta of uremic wild-type and PKCα^−/−^ mice.

All blood vessels were fixed, dehydrated and paraffin-embedded for μCT scanning^29,30^. In both genotypes, the aortic arch exhibited more calcification than the abdominal aorta (Figure 3A-E). Calcification was generally localized to the arterial branches of the abdominal aorta (Figure 3A) or the ascending aorta region of the aortic arch (Figure 3B). The calcifications were either located alongside the elastic fibers (Figure 4A) or across the width of the arterial wall (Figure 4B).

**Figure 3.**
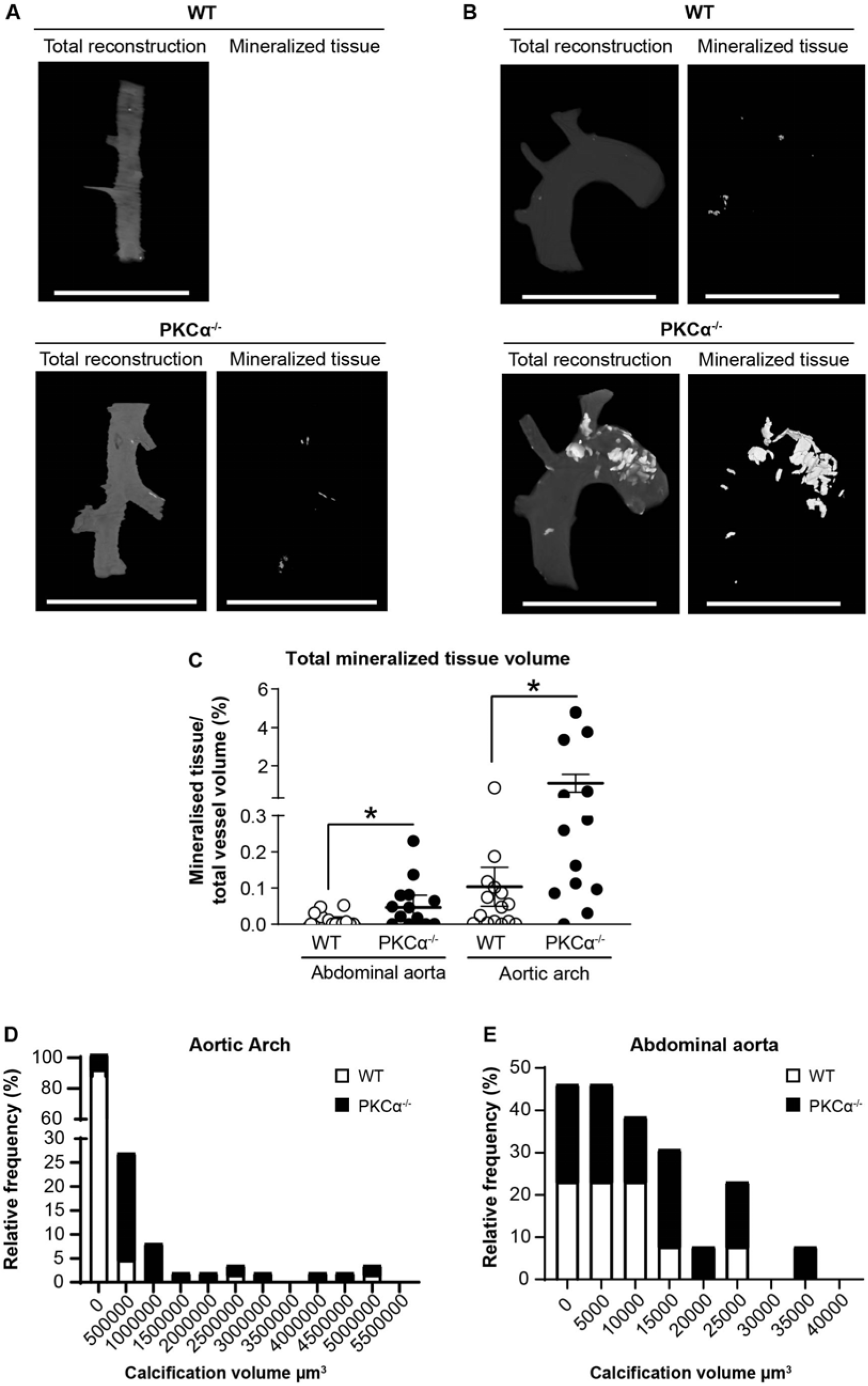
Arterial medial calcification is increased in 5/6 nephrectomy and high phosphate diet fed PKCα^−/−^ mice. Whole vessel and mineralized tissue 3-D reconstructions of representative wild-type (WT) and PKCα^−/−^ (**A**) abdominal aortas and (**B**) aortic arches following 5/6 nephrectomy and high phosphate diet feeding. Blood vessels were analyzed by μCT using a 4x objective with a voxel size between 3.4 μm and 3.8 μm. Scale bar=1500 μm. (**C**) Total mineralized tissue volume expressed as a percentage of total vessel volume (WT n=15; PKCα^−/−^ n=13). Data expressed as median ± interquartile ranges and were analyzed using a Mann Whitney t-test. *P<0.05. (**D, E**) Frequency distribution of individual calcification volumes in the (**D**) abdominal aorta and (**E**) aortic arch from 5/6 nephrectomy and high phosphate diet fed WT and PKCα^−/−^ mice.

**Figure 4.**
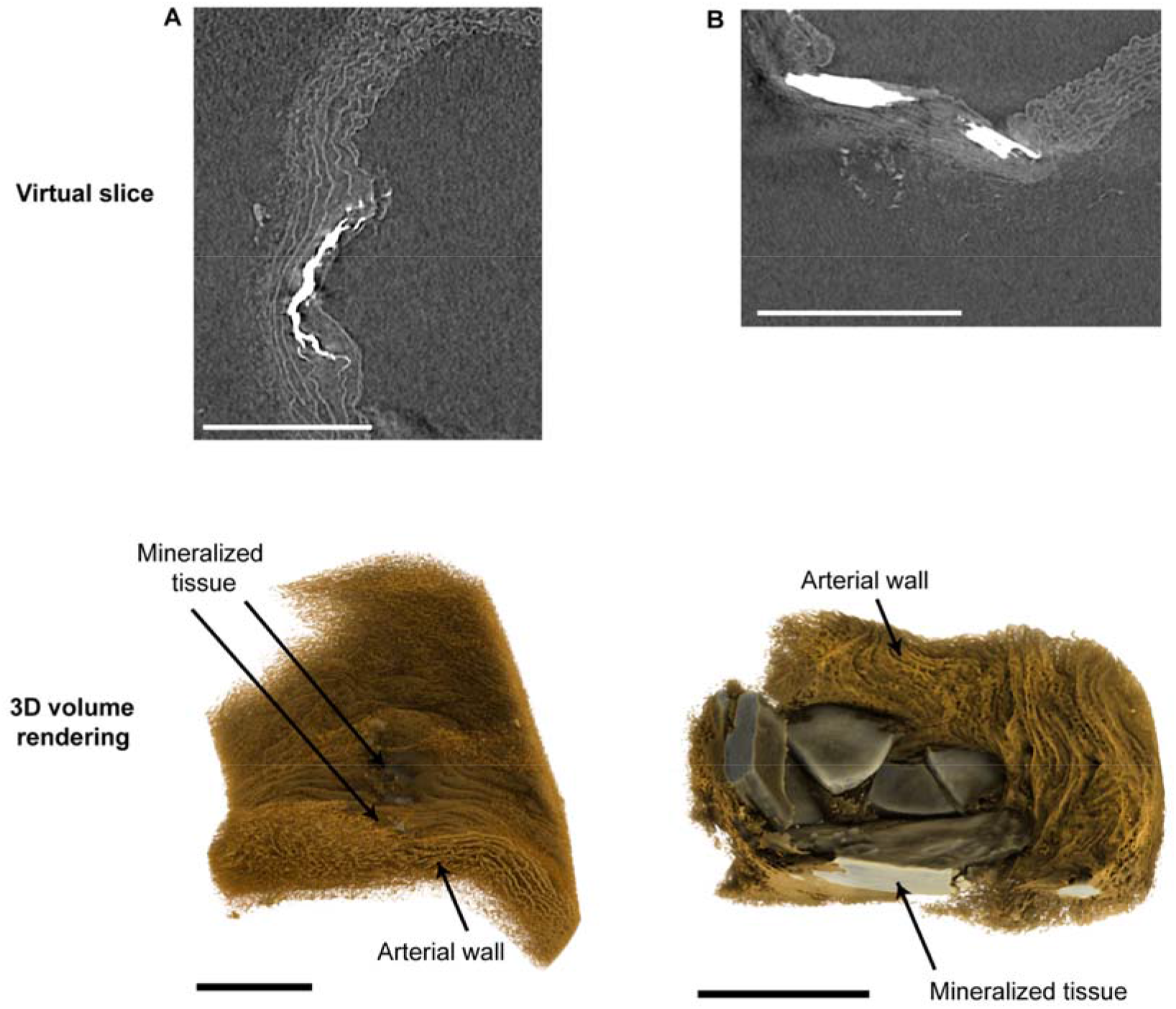
Localisation of arterial medial calcification in mouse blood vessels. Micro-CT phase virtual slices and 3-D reconstructions of aortic arches from 5/6 nephrectomy and high phosphate diet fed PKCα^−/−^ mice using a 20x objective with a voxel size between 0.66 μm and 0.7 μm. Mineralized tissue localizes (A) alongside the elastic fibers within the arterial wall, or (B) across the whole width of the arterial wall. All scale bars=200 μm.

Loss of PKCα increased uremia-induced arterial medial calcification in the abdominal aorta (4.5-fold, P<0.05; Figure 3A&C) and aortic arch (10-fold, P<0.05; Figure 3B&C) when compared to wild-type controls. The scanned blood vessels were subsequently sectioned and stained with von Kossa to confirm calcification; there was a strong correlation (r=0.888; P=0.0003) between the calcification quantified by μCT and von Kossa (Figure S1A). Good agreement between the calcification quantified by the μCT and von Kossa methods was also confirmed using Bland-Altman analysis (Figure S1B). Comparing the frequency distributions of individual calcification sizes (i.e. individual volumes) from each animal, there was no difference in individual calcification size in the abdominal aorta between wild-type and PKCα^−/−^ mice (Figure 3D). However, there was a clear shift towards increased calcification size in the aortic arch of PKCα^−/−^ mice when compared to wild-types (Figure 3E). As there was no correlation between BUN levels and the total volume of calcification in either the abdominal aorta (Figure S2A: r=0.42; P=0.155) or aortic arch (Figure S2B: r=0.26; P=0.384) of PKCα^−/−^ mice, loss of PKCα appears to increase uremia-induced arterial medial calcification independent of its effects on renal function in these mice.

Plasma phosphate concentrations and arterial medial calcification was also compared in 5/6 nephrectomy and high phosphate-fed wild-type and PKCα^−/−^ mice. At a given plasma phosphate concentration, PKCα^−/−^ mice tended to have more calcification in the aortic arch (Figure S3A) and abdominal aorta (Figure S3B) when compared to wild-type controls. There was a trend for plasma phosphate concentration and aortic arch calcification to be correlated in PKCα^−/−^ mice, although this did not reach statistical significance in this study (r=0.539; P=0.06).

### PKCα regulates TGF-β/SMAD2 signaling *in vitro*

We have shown previously that inhibiting TGF-β signaling with SB431542 prevents the increased mineralization observed in PKCα-siRNA treated VSMCs^20^. Therefore, to determine the mechanism by which loss of PKCα increases uremia-induced arterial medial calcification, we further examined the relationship between loss of PKCα and TGF-β signaling *in vitro* and *in vivo*.

Knock-down of PKCα in VSMCs using siRNA^20^ increased TGF-β1-induced SMAD2 phosphorylation when compared to control siRNA-treated cells (P<0.05; Figure 5A). In contrast, pharmacological activation of PKCα using prostratin (10 μM) decreased TGF-β1-induced SMAD2 phosphorylation in VSMCs *in vitro* (P<0.05; Figure 5B). Confirmation that prostratin activated PKCα in these cells was confirmed by immunoblotting for phosphorylated MARK2, a downstream target of PKCα (Figure 5B).

**Figure 5.**
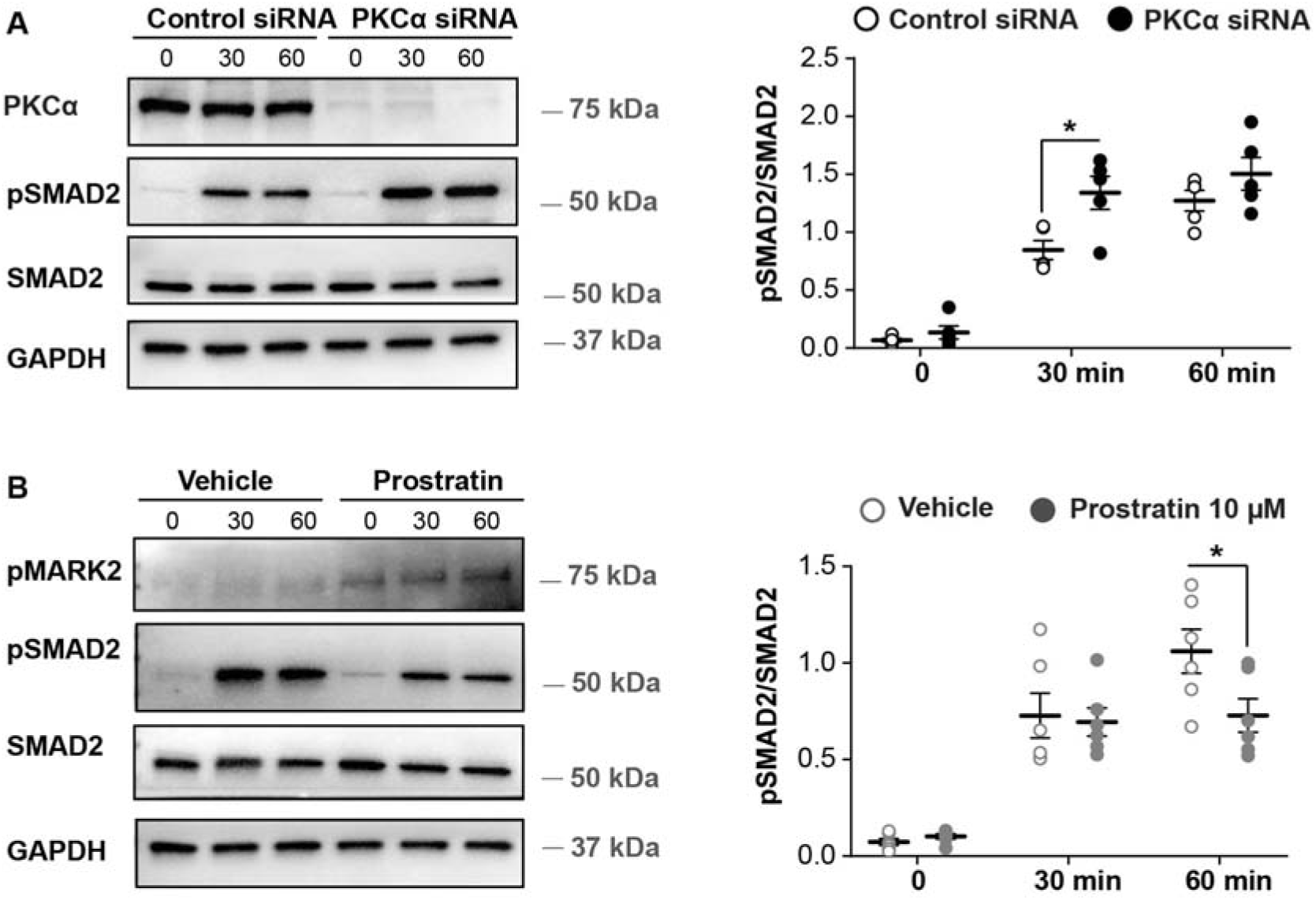
PKCα modulates TGF-β/pSMAD2 signaling in VSMC. (**A**) Control and PKCα siRNA-treated VSMCs were serum-starved for 2□h (0) and stimulated with 0.5 ng/ml TGF-β1 for 30 or 60□min. Knockdown was confirmed by immunoblotting for PKCα. (**B**) VSMCs were serum-starved for 2.5□h (‘0’) with vehicle or prostratin (10□μM) and stimulated with 0.5□ng/mL TGF-β1 for 30 or 60□min. Activation of PKCα signaling with prostratin was confirmed by immunoblotting for phosphorylated MARK2 (pMARK2). (**A, B**) Cell lysates were immunoblotted for phosphorylated SMAD2 (pSMAD2) and total SMAD2; GAPDH was the loading control. Molecular weight markers and the pSMAD2/SMAD2 ratio are shown (n□=□5-6 independent experiments). Data expressed as mean ± SEM and were analyzed using a 2-way ANOVA with Sidak post-hoc tests. *P<0.05.

### Phosphorylated SMAD2 and Runx2 are detected in calcified aortic arches from PKCα^−/−^ mice

To determine whether pro-calcific TGF-β/SMAD2 signaling is also increased in calcified aortic arches from PKCα^−/−^ mice, immunohistochemistry was performed. Phosphorylated SMAD2 immunostaining was detected at sites of calcification in aortic arches from PKCα^−/−^ mice (Figure 6). In contrast, phosphorylated SMAD2 immunostaining was not detected in calcified wild-type arches (Figure 6).

**Figure 6.**
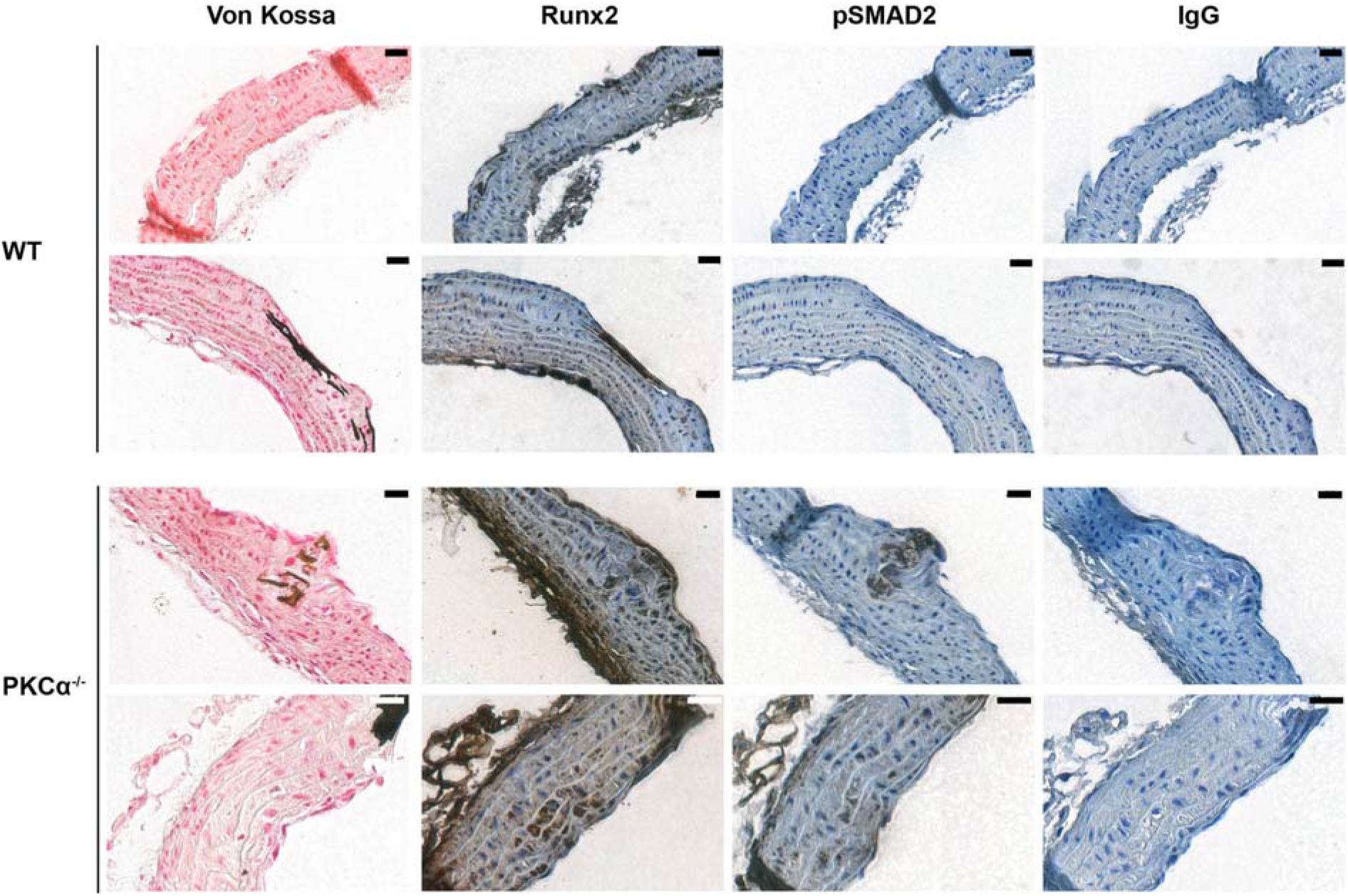
Phosphorylated SMAD2 is detected in calcified aortic arches from 5/6 nephrectomy and high phosphate diet fed PKCα^−/−^ mice. Immunohistochemistry for phosphorylated SMAD2 (pSMAD2) and Runx2 in representative aortic arches from 5/6 nephrectomy and high phosphate diet fed wild-type (WT) and PKCα^−/−^ mice; images from 2 different animals per genotype are shown. Positive antibody staining is shown in brown. Von Kossa staining of adjacent sections and IgG controls are shown. No staining was observed in the IgG controls. Scale bar=20μm.

The osteogenic reprogramming of VSMCs, which is preceded by *de novo* expression of Runx2^17,18^, is critical to the development of vascular calcification in both atherosclerosis and CKD^31,32^. Whilst the osteogenic marker, Runx2, was detected in aortic arches from both WT and PKCα^−/−^ mice, Runx2 staining was more intense and also localized to areas of phosphorylated SMAD2 immunostaining and von Kossa staining in calcified aortic arches from PKCα^−/−^ mice (Figure 6). Together, these studies suggest that loss of PKCα is associated with increased pro-calcific TGF-β/SMAD2 signaling and osteogenic marker expression in calcified aortas.

## Discussion

This is the first demonstration that loss of PKCα increases uremia-induced arterial medial calcification *in vivo*, most likely via a TGF-β/SMAD2 mechanism. We show that TGF-β1-induced SMAD2 phosphorylation is increased in PKCα siRNA-treated VSMCs and that extensive phosphorylated SMAD2 expression is detected in calcified aortic arches from PKCα^−/−^ mice. As we have previously shown that inhibiting TGF-β receptor signaling with SB431542 prevents the increase in mineralization observed in PKCα siRNA-treated VSMCs *in vitro*^20^, these data suggest that loss of PKCα increases calcification both *in vitro*^20,23^ and *in vivo* by up-regulating TGF-β/SMAD2 signaling in VSMCs.

Consistent with our data, previous studies have shown that TGF-β1-induced Smad2 phosphorylation is increased in PKCα^−/−^ podocytes compared to PKCα^+/+^ controls^34^. It has been speculated that PKCα is a key mediator of receptor and membrane endocytosis as there is less internalization of the TGF-β type-1-receptor in PKCα^−/−^ podocytes^34^. It is possible, therefore, that PKCα/TGF-β cross-talk in VSMCs may also be mediated by internalization of TGF-β type-1-receptor.

Whilst our results strongly suggest an important role for TGF-β/SMAD2 signaling in mediating the effects of PKCα in vascular calcification, other signaling pathways may also be involved. For example, WNT/β-catenin signaling promotes vascular calcification^35^ and activation of this signaling pathway is enhanced in the bones of PKCα^−/−^ mice^36^. TGF-β1 also activates the WNT/β-catenin pathway, which in turn stabilizes the TGF-β/SMAD response^37,38^. Loss of PKCα may, therefore, lead to increased pro-calcific TGF-β1/SMAD2 signaling in VSMCs, activating WNT/β-catenin signaling and further promoting vascular calcification. Future studies could determine whether WNT/β-catenin regulates PKCα and TGF-β cross-talk during vascular calcification.

Vascular calcification is thought to be preceded by, or to occur simultaneously, with the osteogenic differentiation of VSMCs^26^. Previous studies have shown that TGF-β induces the expression of Runx2 in a mouse pluripotent mesenchymal precursor cell line^39,40^, and we show that Runx2 localizes to areas of phosphorylated SMAD2 expression in calcified aortic arches from PKCα^−/−^ mice. In contrast, phosphorylated Smad2 was not detected with Runx2 expression in either non-calcified and calcified aortic arches from wildtype mice, suggesting that TGF-β/SMAD2 signaling is not driving vascular calcification in wild-type animals and that a ‘second hit’ is required. Indeed, Runx2 expression is regulated by various mechanisms in VSMCs, including WNT/β-catenin signaling and micro RNAs^41,42^.

Hyperphosphatemia and hypercalcemia are associated with vascular calcification in CKD^8^. Interestingly, calcification occurred in both wild-type and PKCα^−/−^ mice despite the apparent ‘normal’ phosphorus and calcium levels detected after 8 weeks on the high phosphate diet. However, previous studies have shown a rise in calcium and phosphate levels at weeks 1-5 following the initiation of high phosphate diet feeding in mice which have undergone a sub-total nephrectomy, which then normalize by week 7-8^26^.

The pattern and localization of calcification detected in this study are consistent with that observed in previous studies using this uremic DBA/2 mouse model of arterial medial calcification^26, 43–45^; that is, calcification was focally distributed and ‘patchy’. In human specimens where larger calcified deposits have been observed, ‘patchy’ or focally distributed calcifications have also been detected^5, 46–48^. Therefore, the similar patterns of calcification observed in both mouse models and humans validates the translational validity to the human condition of advanced CKD.

Huesa et al.^28^ first reported that μCT imaging could be utilized to visualize and quantify arterial medial calcification in formalin-fixed mouse aortae. Consistent with our study, Huesa et al.^28^ reported that calcification was observed within the ascending aorta region of the aortic arch. However, a limitation of their study was that blood vessels were immersed in corn oil for scanning, precluding subsequent histological analysis of the samples. We modified this protocol based on Walton et al.^30^ and López-Guimet et al.^29^ by embedding mouse blood vessels in paraffin wax, and show that the complete 3-D reconstruction and the accurate quantification of arterial medial calcification by μCT can be combined with subsequent 2-D histological and immunohistochemistry analysis to obtain complementary information about calcification volume, calcification load and signaling mechanisms within the same arterial segment. Furthermore, the high resolution μCT (~0.7 μm) capabilities achieved in this study have enabled the visualization of calcified deposits throughout the blood vessel wall extracellular matrix at a micrometer-scale resolution. This protocol offers several advantages over the *o*-Cresolphthalein assay and histology. Firstly, the *o*-Cresolphthalein assay only quantifies calcium content, and does not visualize the localization or distribution of calcification throughout the blood vessel. Secondly, histological analysis is highly dependent upon *which* sections are stained as calcification is focally localized, and calcification can frequently ‘drop out’ of the tissue when sectioned which precludes the accurate quantification of calcification. The information obtained from a single animal is thus maximized using μCT, which could lead to reductions in the number of animals used in such studies.

Previous studies have shown that PKCα gene expression is *reduced* in kidneys from patients with diabetic nephropathy when compared to non-diabetic controls^50^ and loss of PKCα reduces renal dysfunction in a mouse model of diabetic nephropathy^23^. Whilst these studies have indicated that PKCα inhibitors could be of therapeutic benefit in renal disease patients^23^, our demonstration that loss of PKCα is associated with increased BUN levels and arterial medial calcification in the 5/6 nephrectomy mouse model suggests that vascular calcification and its devastating consequences could be increased if renal disease patients are treated with PKCα inhibitors.

In conclusion, we demonstrate herein that loss of PKCα increases uremia-induced arterial medial calcification by enhancing TGF-β/SMAD2 signaling. Furthermore, pro-calcific TGF-β1-induced SMAD2 phosphorylation is reduced in VSMCs treated with a pharmacological PKCα activator. Further studies are now required to determine whether increasing PKCα activity will prevent or reduce uremia-induced arterial medial calcification in CKD; thus offering a novel therapeutic approach for this devastating pathology.

## Methods

Detailed methods are provided in the Supplementary material.

### Reagents

Reagents were analytical grade and obtained from Sigma-Aldrich (UK) unless otherwise stated. Recombinant human TGF-β1 (#240-B) was from R&D Systems (UK) and used at a final concentration of 0.5 ng/ml. Prostratin (#P0077) was from Sigma-Aldrich (UK) and used at a final concentration of 10 μM; an equivalent volume of vehicle (DMSO) was the control. Immunoblotting antibodies were: phosphorylated SMAD2 (#3108), SMAD2 (#5339) and PKCα (#2056) from Cell Signaling Technology (USA), phosphorylated MARK2 (#34751) from Abcam (UK), GAPDH (60004-1g) and PKCα (21991-1-AP) from Proteintech (UK). Immunohistochemistry antibodies were: phosphorylated SMAD2 (#44-244G) from Thermo Fisher Scientific (UK) and Runx2 (#MAB2006) from R&D Systems (UK).

### Ethical approval

Experiments involving animals were performed in accordance with the UK Animals (Scientific Procedures) Act 1986 (project license number P217A25EF) and received local approval from the University of Manchester Animal Welfare and Ethical Review Board. All efforts were made to minimize suffering throughout the duration of the study. Decisions to cull animals before the end of the study were based on humane end points defined in the project licence.

### Subtotal nephrectomy and high phosphate diet

PKCα^−/−^ mice were generated on the calcification-prone DBA/2 background using CRISPR/Cas9 (Table S1). Wild-type DBA/2 mice were used as controls. Animals had access to a standard chow diet (RM1; Special Diet Services, UK) and water *ad libitum* at all times, and were held in a 12 h-12 h light-dark cycle.

Eleven to twelve week-old female wild-type DBA/2 and PKCα^−/−^ DBA/2 mice underwent a 5/6 nephrectomy in a two-stage procedure to induce uremia and were fed a standard chow diet (RM1) with 1.5% phosphate (Special Diet Services, UK) for 8 weeks^23^. At the end of the study, mice were euthanized with CO_2_. Following excision of the lungs, blood was allowed to pool in the thoracic cavity from which it was collected prior to perfusion of tissues with 10% (v/v) neutral buffered formalin by intraventricular (left ventricle) injection. The aortic arch and abdominal aortae were dissected, removing fat and connective tissue. Tissues were fixed overnight in 10% (v/v) neutral buffered formalin at 4°C.

### Plasma analysis

Blood urea nitrogen (BUN) levels were determined using a colorimetric detection kit (K024-H1; Arbor Assays, USA). Plasma phosphate and calcium concentrations were analyzed on a Roche Cobas 8000 analyzer system using a c702 module.

### Micro-computed tomography (μCT) scans & analysis

Aortic arches and abdominal aortas were dehydrated and paraffin wax-embedded using a Microm STP 120 processor and Microm EC 350-1/2 embedder. Excess wax around the blood vessels was trimmed using a straight-edged blade, and a 25G needle was inserted into the wax so that the vessel axis could be mounted perpendicular to the X-ray source.

All blood vessels were imaged in the Henry Moseley X-ray Imaging Facility (University of Manchester, UK) within the Henry Royce Institute using a Carl Zeiss Versa XRM-520 system (Carl Zeiss: USA) with the X-ray source voltage and power set to 80 kV and 7W, respectively. A low resolution data collection scan of the complete artery cross-section was performed using the 4x objective; for some samples, a second higher resolution region-of-interest scan was performed at a 20x objective. Detector and source distances, exposure times, voxel sizes and scan times are provided in the Online Supplement. The 3D data sets were reconstructed from the original projection data using Zeiss’ Scout-and-Scan™ Reconstructor and scan data were analysed using Avizo 9.7.0 software as described in the Online Supplement.

### Histology

Following μCT scanning, aortic arches were re-embedded in paraffin wax for sectioning. Six-micron sections were cut using a Microm HM 355S and dried overnight at 37°C. Tissue sections, every 60 microns, were analyzed for calcification by staining with von Kossa and counterstaining with nuclear fast red^20^. Images were acquired on a 3D-Histech Pannoramic-250 microscope slide-scanner using a 40x/0.95 *Plan Apochromat* objective (Zeiss). Snapshots of the slide-scans were taken using the Case Viewer software (3D-Histech). ImageJ (version 1.52a) was used to quantify qualification as a percentage of the tissue area.

### Immunohistochemistry

Aortic arches from wild-type and PKCα^−/−^ mice were used for the detection of Runx2 and phosphorylated SMAD2 by immunohistochemistry. Images were acquired on a 3D-Histech Pannoramic-250 microscope slide-scanner as described elsewhere^20^.

### Small interfering-RNA (siRNA)

VSMCs were transfected with siRNA against PKCα (SI01965138, Qiagen, UK) using RNAiMAX (Invitrogen™, Life Technologies, UK). A random control siRNA (#1027281; Qiagen, UK) was used as the control. VSMCs were cultured for 7□days with repeated siRNA transfections every 48-72 h^20^. We previously reported that this PKCα siRNA achieves a 94% and 92% knock-down efficiency in VSMCs at the mRNA and protein level, respectively^20^.

### Immunoblotting

VSMC lysates were analyzed for PKCα (Proteintech 21991-1-AP), phosphorylated MARK2, phosphorylated SMAD2 and total SMAD2 by immunoblotting as described previously (Table S2)^20^. Tissue lysates were analysed for PKCα (Cell Signaling #2056). GAPDH was the loading control. Immunoblots were quantified using ImageJ.

### Statistical analysis

Data with a normal distribution are presented as mean□±Cstandard error of the mean (SEM); data without a normal distribution are presented as median ±□interquartile range. Differences in survival were determined using a Gehan-Breslow-Wilcoxon test. Data were analyzed using a Student’s t-test, or a Mann-Whitney test if a normal distribution was not achieved. Data with two or more variables were analyzed using a Kruskall-Wallis test or 2-way ANOVA with Tukey post-hoc comparisons. A Spearman test or Bland-Altman analysis was performed to analyze correlation and agreement between data sets, respectively. All analysis was performed using GraphPad Prism software (California, USA); *P*<0.05 was considered statistically significant.

### Disclosure

None

## Acknowledgements

We thank the Biological Service Facility (University of Manchester, UK) for their support and care of all the animals used in this study; Maj Simonson-Jackson in the Genome Editing Unit (University of Manchester, UK) for help in generating the PKCα^−/−^ mice; Roger Meadows (University of Manchester, UK) for help with bioimaging.

