## Supplemental Methods for "Loss of PKCα increases arterial medial calcification in a uremic mouse model of chronic kidney disease"

**Supplementary Methods**

**Generation of PKCα^-/-^ mice**

sgRNA targeting Exon 3 of the PKCα gene were selected that were predicted to have no off-target sites of less than 3 mismatches elsewhere in the genome (sgRNA1 gttcgttacgttctcttgtc-*cgg*, sgRNA 2 gatgttaactcgaatgtctg-*agg* (Figure 1A). The gRNA, delivered as alt-R crRNA combined with tracrRNA (Integrated DNA Technologies, USA) and Cas9 protein (New England Biolabs, UK) were microinjected into the pronuclei of DBA/2 mouse zygotes using standard techniques, with the modification of using older females (8-12 weeks), which enhanced DBA/2 superovulation and thus the embryo yield. A linear dsDNA donor template, comprising the floxed exon with 800bp homology arms, was also included to attempt to generate loxP flanked conditional allele but we detected no knock in by Homology Directed Repair. Post-injection, surviving embryos were transferred to pseoudopregnant CD1 females. After genomic DNA extraction from ear punches pups were genotyped by amplifying the target region with the following primers: PKCα forward 5’-cttttctctcctcagtttgctgttt-3’, PKCα reverse 5'-gacgaagtgagaaaaccgagat-3’. PCR conditions were: 94°C for 10 min followed by 37 cycles of 94°C, 20 sec; 60°C, 20 sec; 72°C, 20 sec and a final 10 min incubation at 72°C. Mice harbouring InDel containing alleles were identified by size change from wild-type amplicon size (230bp) using Qiaxcel analysis (Qiagen, UK) (Figure 1B). Three mice (founders 1, 8 and 10) were taken forward for Sanger sequencing and changes predicted to lead to frameshift mutations in the exon, and thus lead to gene knock-out, identified (Figure 1C).

PKCα founder 8 did not produce offspring, and PKCα founder 10 did not produce offspring containing the frameshift mutation. PKCα founder 1 was bred with WT DBA/2 to establish a colony, and bred to homozygosity. The absence of PKCα in these mice was confirmed by immunoblotting of heart and kidney lysates (Figure 1D). Animals had access to a standard chow diet (RM1; Special Diet Services, UK) and water ad libitum at all times, and were held in a 12 h–12 h light–dark cycle.

**Subtotal nephrectomy and high phosphate diet**

All surgery was performed under isoflurane anesthesia (4% in oxygen at 2 L/min) with subcutaneous administration of fluids (0.2 mL saline) and buprenorphine (0.003 mg) at the beginning of each surgical procedure.

Wild-type and PKCα^-/-^ mice underwent a 2/3^rd^ resection of the left kidney at week 0: the lower and upper kidney poles were removed and tissue glue (Histoacryl, Braun, UK) was applied to curtail bleeding. Two weeks later (week 2), a total right nephrectomy was performed. To aid recovery, mice were kept in a warming cabinet for at least one night following surgery 2. Percentage renal mass removed was calculated as described^1^. On average, 65.1 ± 3.8% (SD, *n*=15) of renal mass was removed from wild-type and 66.4 ± 2.9% (SD, *n*=13) from PKCα^-/-^ mice, with no statistically significant difference between groups. One week after the 2^nd^ surgery, mice were placed on a standard chow diet (RM1) with 1.5% phosphate (Special Diet Services, UK) for 8 weeks. The phosphate was increased using di-sodium hydrogen phosphate; thus, the sodium concentration of the 1.5% phosphate was increased from 0.23% to 1.73%. The high phosphate diet was provided *ad libitum* as both pellets and fresh mash daily. Hydrogel was also provided to the mice *ad libitum*. Mice were checked and weighed at least 3 times a week for the duration of the study.

**Micro-computed tomography (µCT) scans**

Aortic arches and abdominal aortas were dehydrated and paraffin wax-embedded using a microm STP 120 processor and Microm EC 350-1/2 embedder. Excess wax was trimmed using a straight-edged blade, and a 25G needle was inserted into the wax so that the vessel axis could be mounted perpendicular to the X-ray source.

All blood vessels were imaged in the Henry Moseley X-ray Imaging Facility (University of Manchester, UK) within the Henry Royce Institute using a Carl Zeiss Versa XRM-520 system (Carl Zeiss: USA) with the X-ray source voltage and power set to 80 kV and 7W, respectively. The detector and source were positioned ~14 mm and ~15 mm respectively from the sample to maximise the field of view and achieve a small amount of X-ray phase contrast. A low resolution data collection scan of the complete artery cross-section was performed using the 4x objective, achieving a voxel size between 3.4 and 3.8 μm. For some samples, a second higher resolution region-of-interest scan was performed at a 20x objective achieving a voxel size between 0.66 and 0.74 μm. Exposure times per radiograph were between 1 and 1.5 sec for the 4x scans, and between 15 and 20 sec for the 20x scan. Exposure times and voxel size were dependent on the sample size. In both cases, each scan consisted of 1601 projections, collected over 360^o^ rotation. Scan times were approximately 1 hour for 4x objective and 12 hours for the 20x objective. The 3D data sets were reconstructed from the original projection data using Zeiss’ Scout-and-Scan™ Reconstructor.

Scan data were analyzed using Avizo 9.7.0 software. A median filter was used to smooth out image noise, before segmenting the blood vessel and mineralized tissue for quantification (Figure 5A). Mineralized tissue was identified and selected using localized thresholding functions; a global thresholding function could not be used due to the presence of X-ray dense material inside the paraffin wax. The low absorption contrast between the vessel and the surrounding wax required manual intervention to segment out the total vessel volume. The segmented samples were rendered in 3-D and the percentage of mineralized tissue expressed as total mineralized tissue volume over the total vessel volume.

**Immunohistochemistry**

Aortic arches from wild-type and PKCα^-/-^ mice were used for the detection of Runx2 and phosphorylated SMAD2 by immunohistochemistry. Endogenous peroxidase activity was blocked with 3% (v/v) H_2_O_2_, and antigen retrieval performed with 1x universal antigen retrieval buffer (Abcam, UK; Runx2) or 100 mM citric acid monohydrate, pH 6 (phosphorylated SMAD2) at 95°C for 20 min. Non-specific binding was blocked with 2% (v/v) rabbit serum (Runx2) or 2% (v/v) goat serum (phosphorylated SMAD2) in phosphate buffered saline for 1 h at room temperature. Antibodies against phosphorylated SMAD2 (1:200) or Runx2 (1:100) were diluted in the same solution used to block and incubated with the tissue at 4°C overnight. Tissue incubated with concentration-matched, non-immune rabbit IgG (Sigma-Aldrich, UK) or rat IgG^2b^ (Bio-Rad, UK) were used as controls. The following day, tissues were incubated with a biotin-conjugated secondary antibody (1:200 dilution in same solution used to block) and positive immunoreactivity was detected using the ABC system (Dako, Denmark) and 3’3’-diaminobenzidine (Sigma-Aldrich, UK); nuclei were counterstained with Harris’s hematoxylin. Images were acquired on a 3D-Histech Pannoramic-250 microscope slide-scanner using a 40x/0.95 Plan Apochromat objective (Zeiss). Snapshots of the slide-scans were taken using the Case Viewer software (3D-Histech).

**Cell culture & small interfering-RNA (siRNA)**

Bovine VSMCs were isolated and cultured as described^2^. Two different preparations of uncloned VSMCs were used for these experiments at passage 11-13. VSMCs were transfected with siRNA against PKCα (SI01965138, Qiagen, UK) using RNAiMAX (Invitrogen™, Life Technologies, UK); effective knock-down of PKCα expression in bovine VSMCs using this siRNA has been reported previously^2^. A random siRNA (#1027281; Qiagen, UK) was the control. Both siRNAs were used at a final concentration of 20 nM. For signaling assays, VSMCs were cultured for 7 days with repeated siRNA transfections every 48–72 h.
