## Supplemental Table 1 for "Loss of PKCα increases arterial medial calcification in a uremic mouse model of chronic kidney disease"

**Table S1. Founder PKCα^-/-^ mice generated by CRISPR/Cas9.**

| **Founder** | **Allele 1** | | **Allele 2** | | **Predicted effect on gene** | **Amplicon (WT=230)** |
| --- | --- | --- | --- | --- | --- | --- |
|  | g399 | g392 | g399 | g392 |  |  |
| 1 | -8bp | 2bp SUB | +1bp | WT | Allele 1 = KO  Allele 2 = KO | 222  231 |
| 8 | -5bp | -4bp | WT | WT | Allele 1 = KO | 221  230 |
| 10 | -8bp | WT | WT | WT | Allele 1 = KO | 222  230 |

WT, wild-type.
