## Supplemental Table 2 for "Loss of PKCα increases arterial medial calcification in a uremic mouse model of chronic kidney disease"

**Table S2. Immunoblotting antibodies**

| **Protein** | **Blocking agent** | **Primary antibody and catalogue number** | **Dilution and duration** | **Secondary antibody** | **Dilution and duration** |
| --- | --- | --- | --- | --- | --- |
| PKCα | 5% (w/v) milk in TBST | Cell Signaling (#2056) | 1:500; o/n 4°C | Dako goat anti-rabbit HRP | 1:1000; 1h 4°C |
| PKCα | 5% (w/v) milk in TBST | Proteintech  (#21991-1-AP) | 1:750; o/n 4°C | Dako goat anti-rabbit HRP | 1:1000; 1h 4°C |
| pSMAD2  (Ser^465/467^) | 5% (w/v) BSA in TBST | Cell Signaling (#3108) | 1:500 or  1:1000; both o/n 4°C | Dako goat anti-rabbit HRP | 1:1000; 1h 4°C |
| SMAD2 | 5% (w/v) BSA in TBST | Cell Signaling (#5339) | 1:1000; o/n 4°C | Dako goat anti-rabbit HRP | 1:1000; 1h 4°C |
| pMARK2  (Thr^595^) | 5% (w/v) BSA in TBST | Abcam  (ab34751) | 1:500; o/n 4°C | Dako goat anti-rabbit HRP | 1:1000; 1h 4°C |
| GAPDH | 5% (w/v) milk in TBST | Proteintech (60004-1g) | 1:2000; o/n 4°C | Dako rabbit anti-mouse HRP | 1:1000; 1h 4°C |

o/n, overnight
