## Supplemental Figures 1,2&3 for "Loss of PKCα increases arterial medial calcification in a uremic mouse model of chronic kidney disease"

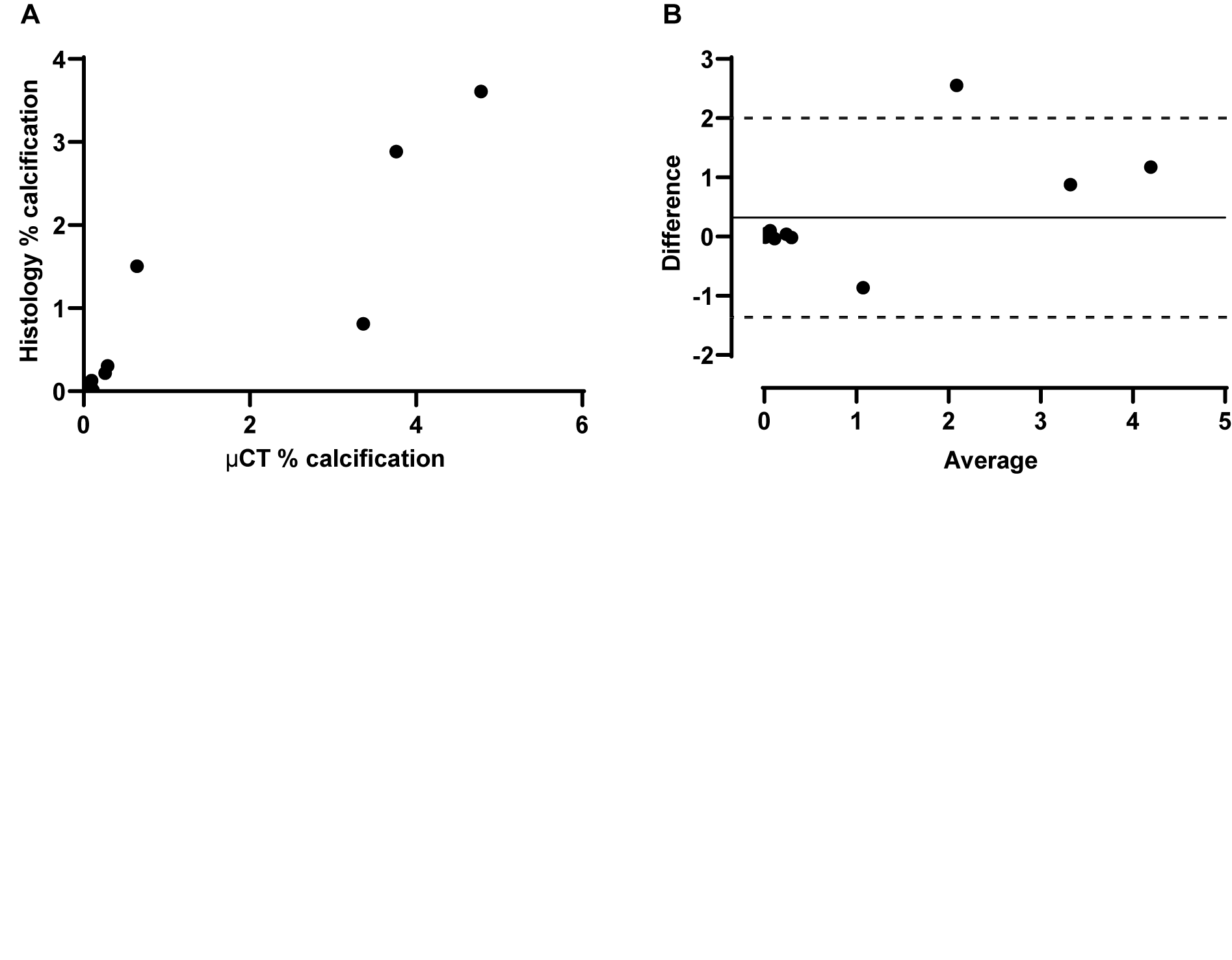

**Figure S1. Correlation between calcification quantification by histology and µCT in 5/6 nephrectomy and high phosphate diet fed wild-type and PKCα^-/-^ mice.** (**A**) Spearman correlation (r=0.888; P=0.0003) and (**B**) Bland-Altman analysis of aortic arch calcification quantified by µCT and von Kossa staining from 5/6 nephrectomy and high phosphate diet fed wild-type and PKCα^-/-^ mice.

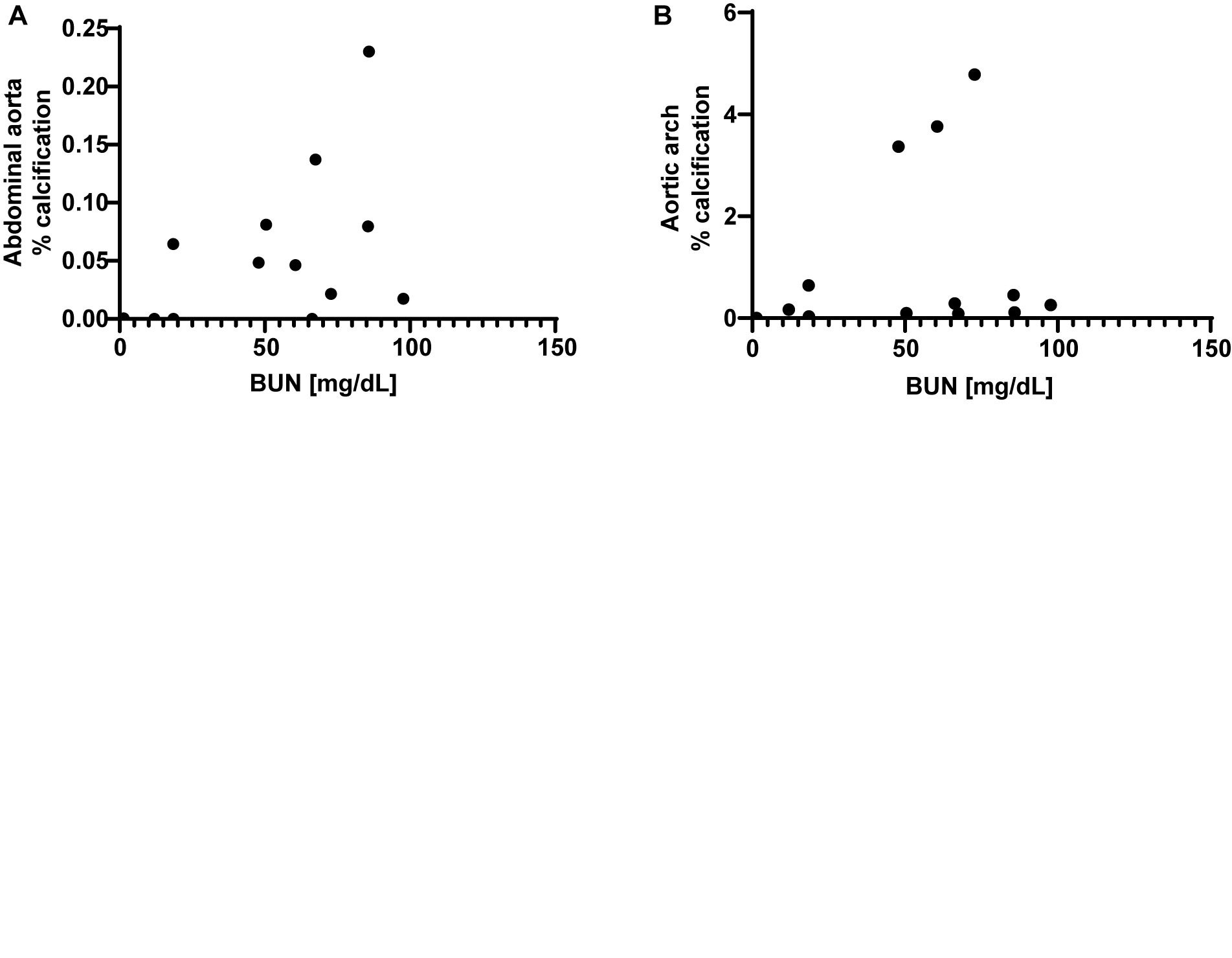

**Figure S2. Correlation between BUN levels and arterial calcification in 5/6 nephrectomy and high phosphate diet fed PKCα^-/-^ mice.** Spearman correlation was performed between plasma BUN levels and arterial calcification in the (**A**) abdominal aorta (r=0.42; P=0.155) and (**B**) aortic arch (r=0.26; P=0.384) from 5/6 nephrectomy and high phosphate diet fed PKCα^-/-^ mice.

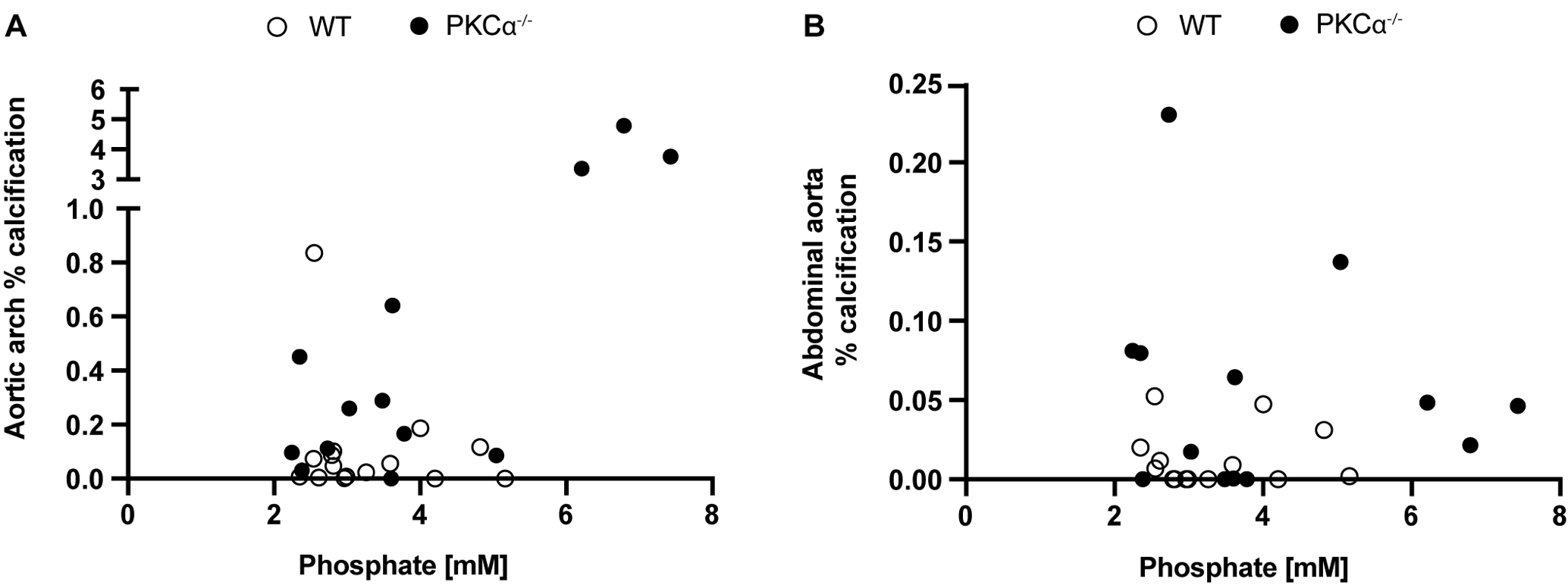

**Figure S3. Correlation between plasma phosphate levels and arterial calcification in 5/6 nephrectomy and high phosphate diet fed wild-type and PKCα^-/-^ mice.** Spearman correlation was performed between plasma phosphate levels measured at the end of the study and arterial calcification in the (**A**) aortic arch and (**B**) abdominal aorta of wild-type (WT) and PKCα^-/-^ mice following 5/6 nephrectomy and high phosphate diet feeding. There was a trend for plasma phosphate concentration and aortic arch calcification to be correlated in PKCα^-/-^ mice (r=0.539; P=0.06).
